# TabSyM: A Generative Pipeline for Small Multi-Cohort Omics Tabular Data

**DOI:** 10.1101/2025.07.14.664738

**Authors:** Nengneng Yu, Yuefan Wang, Lindsey Kathleen Olsen, Bing Zhang, Hui Zhang, Zaoxing Liu

**Affiliations:** Department of Computer Science, University of Maryland, College Park, MD, USA; Department of Pathology, Johns Hopkins University School of Medicine, Baltimore, MD, USA; Lester and Sue Smith Breast Center, Baylor College of Medicine, Houston, TX, USA; Department of Molecular and Human Genetics, Baylor College of Medicine, Houston, TX, USA

**Keywords:** Omics, GenAI, Synthetic Data, Tabular Data

## Abstract

Machine learning applications in biomedicine such as omics data analysis are frequently hindered by datasets that are small, high-dimensional, and affected by batch effects across different patient cohorts. To address these challenges, we introduce TabSyM, a modular generative pipeline that synthesizes high-quality, task-relevant data to improve predictive modeling. TabSyM integrates three key stages: it extends a diffusion-based model (TabDDPM) to generate new omics data, employs a novel task-aware sampling mechanism guided by Bayesian optimization to select the most informative synthetic samples, and uses a Multi-Domain Adversarial Network (MDAN) to align data distributions for cross-cohort generalization. We validated our pipeline on a challenging, real-world task of predicting 3-year survival in gastric cancer patients from high-dimensional scRNA-seq data across five cohorts. The full TabSyM pipeline achieved a 30.2% AUROC improvement over the best tree-based models and an 11.5% AUROC gain over leading automated machine learning frameworks. Furthermore, the generative and sampling components are model-agnostic and can substantially boost the performance of classical models like XGBoost independently. These results establish that combining generative modeling with task-aware sampling and domain adaptation provides a robust and effective strategy for overcoming critical data limitations in biomedical tabular data analysis.

## Introduction

Machine learning is playing an essential role in analyzing complex multi-omics data such as genomics, transcriptomics, proteomics, and metabolomics ^1 2 3 4^. However, realizing its full potential remains difficult due to fundamental data limitations. First, acquiring high-throughput molecular profiles is both costly and logistically challenging, often resulting in small patient cohorts—typically only dozens to hundreds of samples. Unlike in domains such as imaging or text data, where the number of training samples often exceeds the number of features, biomedical datasets are typically high-dimensional with far more features than samples ^5 6^. This imbalance hinders effective model training and generalization. Additionally, integrating data across studies introduces technical variability, or batch effects, alongside biological heterogeneity across cohorts ^7^. These issues collectively compromise model robustness and generalizability, ultimately limiting the development of reliable predictive tools and impeding reproducible scientific discovery in biomedical settings.

To address data scarcity and high dimensionality in leveraging machine learning for multi-cohort and multi-omics data analysis, traditional strategies have primarily focused on either reducing feature space or increasing sample size through cohort aggregation. Feature engineering and dimensionality reduction methods, such as principal component analysis (PCA) ^8^, autoencoders ^9^, aim to compress high-dimensional inputs into lower-dimensional representations ^10^. While such approaches can reduce overfitting in small-sample settings, important biological signals risk being discarded during compression ^11^ and they are often insufficient when the number of samples is extremely limited ^12^. Alternatively, large-scale data aggregation efforts, such as biobank ^13 14^-scale multi-institutional datasets, attempt to overcome sample size limitations by pooling data across studies. However, these large datasets are difficult to access, require significant coordination across sites, and often obscure fine-grained biological variation, especially when focusing on specific subtypes or rare conditions ^15^. As a result, these traditional responses remain inadequate for enabling robust and generalizable learning in many real-world biomedical settings ^6 16^.

Data augmentation represents another approach aimed to alleviate data scarcity by expanding the training set with synthetic samples ^5 17^. The simplest approach, random oversampling (ROS) ^18^, duplicates existing minority class samples to balance class distributions. While effective in preventing classifiers from being biased toward the majority class, ROS does not introduce any new variation and increases the risk of overfitting, particularly in low-sample biomedical settings ^19 20^. To address this, interpolation-based methods, such as SMOTE ^21^ and its variants, generate convex combinations of existing samples in a computationally inexpensive manner. However, these techniques still operate without regard for the underlying data manifold, which is particularly problematic in biomedical omics data where class boundaries are complex and biologically entangled ^22 23 24 25^. As a result, both replication-based and interpolation-based oversampling techniques often fail to capture meaningful biological variability and may introduce noise or distort class structure.

In recent years, deep generative models have been widely adopted for data augmentation across various machine learning domains. Methods such as generative adversarial networks (GANs) ^26 24^, variational autoencoders (VAEs) ^27^, and diffusion models ^28^ have achieved notable success in domains such as vision ^29^, language ^30^, and, more recently, network ^31 32^ and systems modeling. These approaches learn complex data distributions from training data, allowing them to generate novel samples that reflect global structure and fine-grained variation. In biomedical research, generative models have been explored for tasks such as drug discovery and biological sequence generation ^33 34^. However, most existing work is narrowly tailored to specific tasks and lacks a generalizable, plug-and-play framework ^25^. Furthermore, a critical limitation remains: generative models typically assume that all synthetic samples are equally informative. Without mechanisms to assess or filter generated data for task relevance, synthetic augmentation can introduce misleading signals and degrade performance ^35^, especially in high-dimensional, low-sample biomedical settings ^36^.

Besides data scarcity, another major challenge in biomedical machine learning is dataset heterogeneity. Biomedical datasets are often collected across different institutions or platforms, introducing significant batch effects and technical variability. A common strategy to address this issue is domain adaptation. Traditional statistical approaches such as ComBat ^37^, SVA ^38^, and RUV ^39^ have been widely adopted in the bioinformatics community, alongside more recent deep learning methods ^40 41^ that explicitly model inter-cohort variation. However, a fundamental limitation remains: domain adaptation methods alone do not generate new training samples. As a result, their effectiveness remains constrained by the size and diversity of the original dataset ^42^.

To address both data scarcity and cohort heterogeneity, we propose TabSyM, a generalizable synthetic data augmentation pipeline that selects task-aware samples and integrates domain adaptation. At its core, our approach synthesizes high-dimensional molecular data using a diffusion-based generative model. Unlike traditional augmentation strategies that indiscriminately incorporate all generated samples, we introduce a task-aware sampling mechanism that selects only the most informative synthetic data for downstream learning. To further enhance generalization across cohorts, we partition data from different cohorts and incorporate synthetic samples as an external source into domain adaptation models. We validate our framework on a real-world biomedical prediction task, demonstrating consistent performance improvements over state-of-the-art methods.

## Results

### TabSyM Overview

To address the challenges posed by limited sample sizes, high-dimensional feature spaces, and cohort heterogeneity in biomedical data, we propose TabSyM, a modular learning pipeline that integrates generative augmentation, task-aware sampling, and multi-source domain adaptation. As illustrated in Figure 1, the framework consists of three key components: a diffusion-based generative model that synthesizes high-dimensional omics data, a sampling mechanism that selects task-relevant synthetic subsets, and a domain adaptation module that aligns multiple cohorts, including both real and synthetic sources, into a unified representation space. Each component is designed to address a specific barrier to robust prediction: generation expands data diversity, sampling promotes relevance, and adaptation improves generalization across datasets. Together, these modules form a cohesive strategy for improving biomedical prediction tasks under low-resource, high-variance conditions.

**Figure 1.**
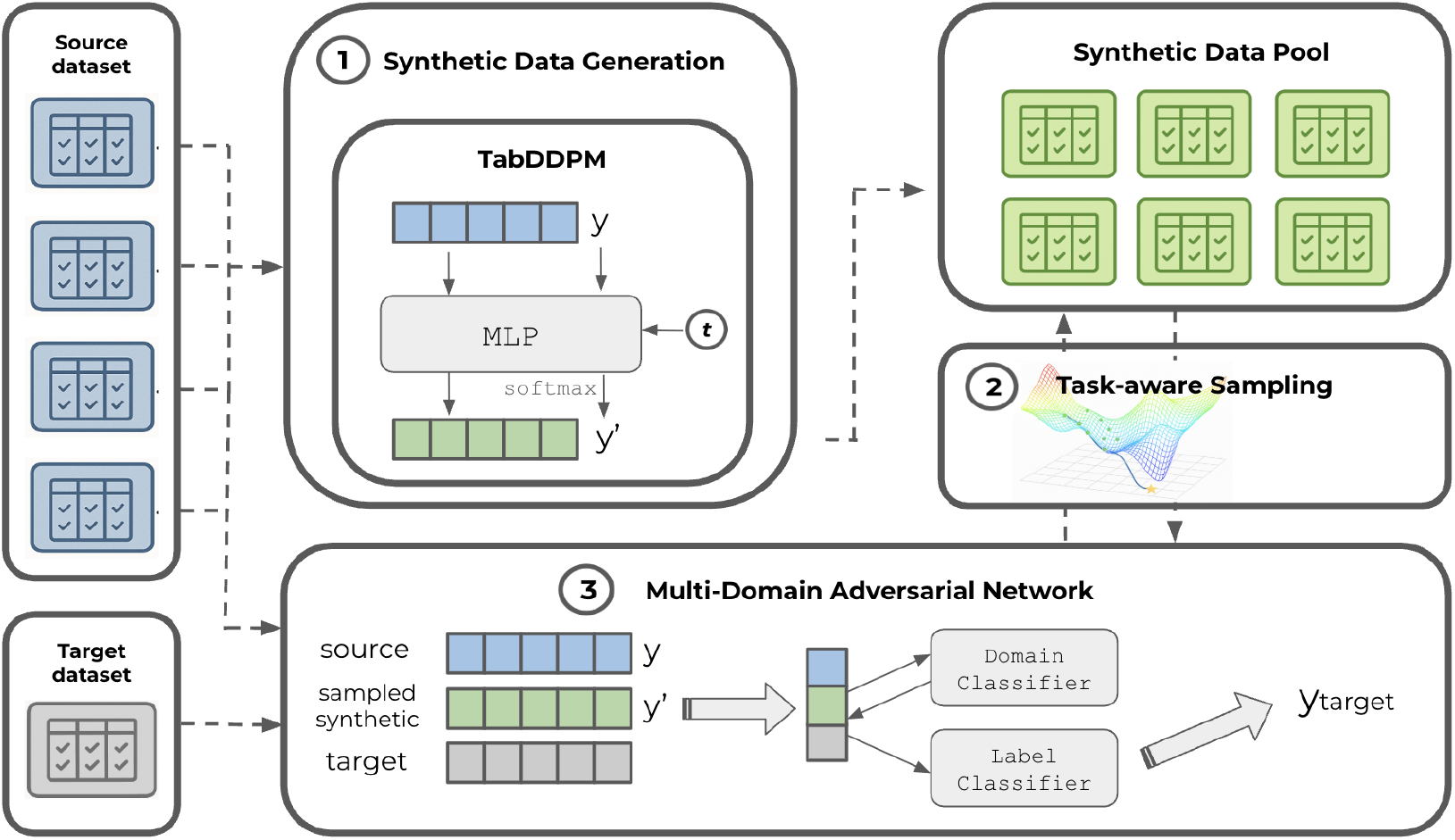
Overview of the TabSyM pipeline. The architecture consists of three stages: (1) synthetic data generation using TabDDPM trained on source datasets, (2) multi-domain adversarial training using both source, target domains and sampled synthetic data, and (3) task-aware sampling to select optimal synthetic samples. The generated synthetic data are pooled and filtered by downstream performance. The MDAN model leverages both real and selected synthetic data to learn domain-invariant features and produces the final prediction for the target domain.

To break down our learning pipeline, we begin by dividing the available data into source datasets and a target dataset. The source datasets may include multiple cohorts, as our model explicitly supports multi-source training. These datasets serve as the foundation for model learning and are assumed to have complete label annotations. In contrast, the target dataset represents the domain of interest for deployment, which may lack comprehensive label information due to real-world constraints. Our pipeline is designed to support this label-sparse setting without sacrificing predictive performance.

The first stage of the pipeline involves synthetic data generation using a diffusion-based generative model - TabDDPM ^43^. We train the TabDDPM solely on source datasets, deliberately excluding target data to minimize the risk of information leakage. This produces diverse sets of synthetic samples that mirror the distributions of the source cohorts.

In the second stage, we integrate the source datasets, their corresponding synthetic data, and the unlabeled target dataset into a Multi-Domain Adversarial Network ^40^ for downstream training and prediction. Each source cohort and its synthetic extension are treated as separate domains, contributing both their input features and label information to MDAN training. The target dataset, on the other hand, is incorporated in an unsupervised manner, contributing only feature representations to enable domain alignment without label usage. This design enables the model to learn domain-invariant features and generalize effectively to the label-scarce target cohort.

During MDAN training, we apply a task-aware sampling strategy to filter the most relevant synthetic data. Given that the volume of generated samples can significantly exceed that of real samples, we use Bayesian Optimization ^44 45^ to automatically select a subset of synthetic data that maximizes downstream task performance. This mechanism ensures that only the most relevant synthetic information is propagated through the adaptation model.

To ensure the robustness of both model training and sampling selection, parameter tuning and sampling evaluation are conducted using a dedicated validation set. Specifically, we reserve a subset of labeled samples from the target dataset as an independent validation set, which is entirely excluded from the training process. All hyperparameter tuning decisions and the objective function for Bayesian Optimization are based solely on performance on this validation set. This separation allows us to maximize generalization performance while avoiding information leakage, and ensures that both the MDAN model and the selected synthetic samples generalize well to the unseen portions of the target domain.

### Synthetic data and Task-aware sampling

High-quality synthetic data serves as the foundation of our pipeline to address the small-sample, high-dimensional nature of tabular biomedical datasets. Ideally, generated samples should preserve complex relationships within the data while also remaining robust to high feature dimensionality. However, many conventional data augmentation approaches fail to meet this bar. Shallow methods such as SMOTE ^21^ or random oversampling either duplicate data points or interpolate them linearly, offering little diversity and risking overfitting ^22^. Deep generative methods that are specifically designed for tabular data like CTABGAN ^46^ and TVAE ^47^ have demonstrated success in other domains but tend to struggle with the sparsity, heterogeneity, and high dimensionality typical of biomedical data ^48^.

To address these limitations, we adopt TabDDPM, a recently proposed denoising diffusion model tailored for tabular data (see Figure 2). TabDDPM ^43^ models numerical and categorical features jointly via Gaussian and multinomial diffusion processes, respectively, and learns to iteratively transform random noise into high-fidelity tabular records (See Method Section). Unlike GANs, which often collapse or fail in sparse feature spaces, TabDDPM utilizes a Gaussian diffusion process for noise addition and removal, enabling it to effectively capture multimodal dependencies and generate diverse, realistic samples. We benchmark TabDDPM against multiple baseline synthesizers, including

**Figure 2.**
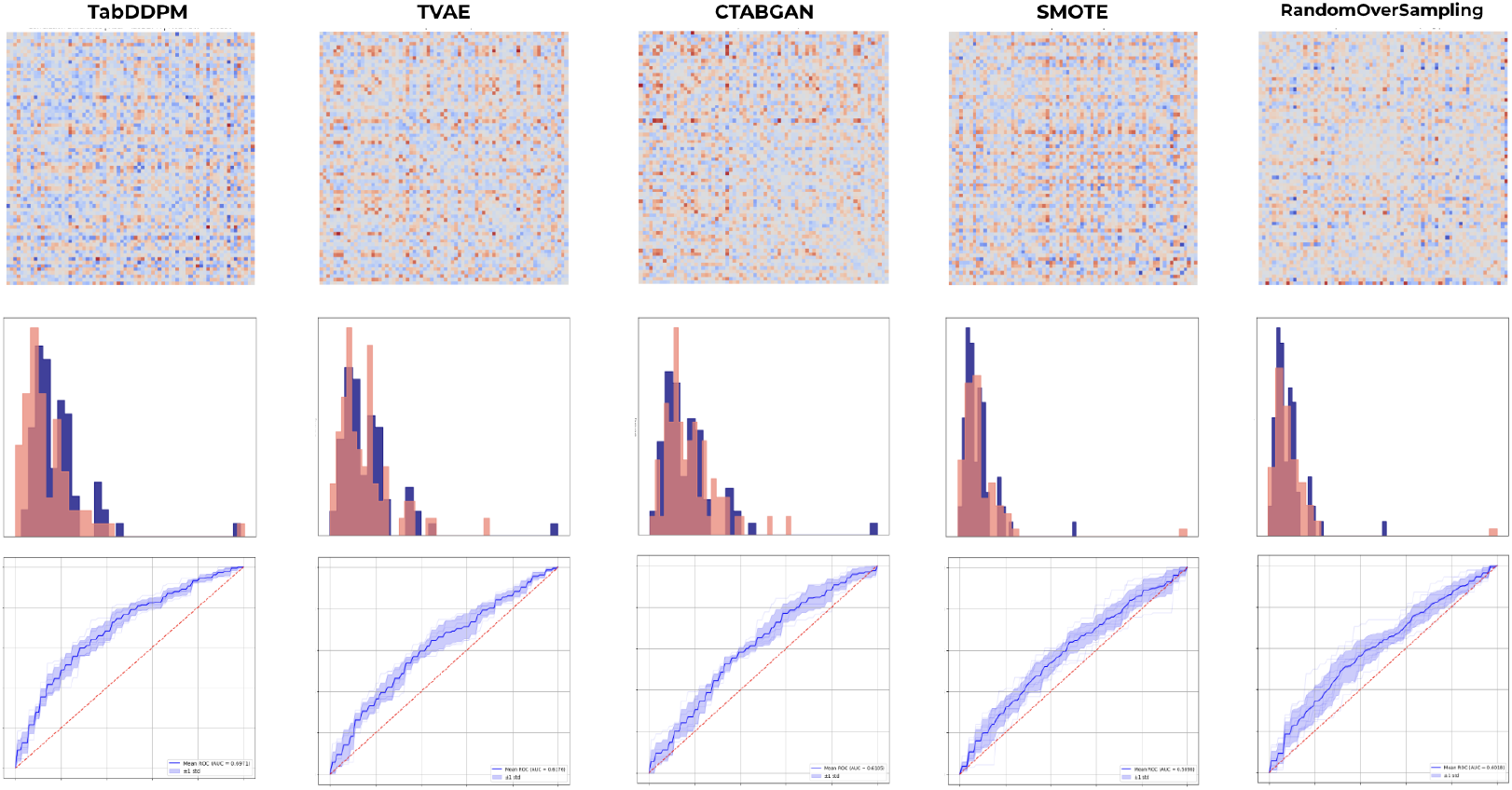
Performance comparison across different synthesizers. The first row shows the difference between the correlation matrices of real and synthetic datasets, with more intense red or blue indicating a larger difference. The second row compares the overall feature value distributions across samples (blue: real data, red: synthetic data). The third row presents the AUROC curves for the final classification task using different synthesizers within our full pipeline.

SMOTE, ROS, CTABGAN, and TVAE. As shown in Figure 2, augmenting our pipeline with TabDDPM consistently leads to stronger downstream predictive performance. Compared to TVAE, our pipeline achieves a significant improvement in both metrics, with an AUROC increase of 22.5% (0.698 vs 0.570) and an F1 gain of 23.3% (0.650 vs 0.527). Against CTABGAN, the performance gap is even more pronounced, with AUROC improving by 18.5% (0.698 vs 0.589) and F1 nearly doubling by 98.2% (0.650 vs 0.328). When compared to ROS, our method improves AUROC by 18.1% (0.698 vs 0.591) and F1 by 11.1% (0.650 vs 0.585). Also, similar performance improvement over SMOTE, we observe an AUROC gain of 14.8% (0.698 vs 0.608) and an F1 improvement of 7.4% (0.650 vs 0.605). These results highlight the overall strength of our selected synthesizer across diverse state-of-the-art baseline synthesizers. In particular, CTABGAN struggles to generalize under high-dimensional settings, often failing to capture complex feature interactions and occasionally generating samples only to one single class. This further emphasizes the importance of choosing the right generative model in biomedical applications.

While synthetic data generation has shown clear benefits particularly in small-sample regimes where training data is limited, the trade-off between the quality and diversity of generated samples remains an open challenge ^49 50^. Prior work has shown that focusing solely on generating samples that closely resemble the original data distribution may inadvertently limit model performance, as such samples often lack the variation needed to support generalization ^51^. This has motivated efforts to identify more selective sampling strategies that go beyond distributional similarity and directly optimize for task relevance. In domains such as computer vision ^51 52^ and language modeling ^53^, task-aware or validation-guided sampling methods have been shown to significantly improve the effectiveness of synthetic data. These findings suggest that selecting the right synthetic samples, rather than simply pick the most realistic-looking ones, is crucial for effective augmentation, particularly in high-dimensional, small-cohort biomedical settings.

To address this, we introduce a task-aware sampling strategy that selects synthetic subgroups based on their estimated utility for the target task. Specifically, for each generated group, we evaluate multiple label-wise sampling ratios and use Bayesian Optimization ^45^ on a held-out validation set to select the top-k sample sets with the highest task performance. These groups are then aggregated via prediction score ensembling. This approach allows us to filter out redundant or uninformative samples while preserving diversity across synthetic subpopulations. As shown in Figure 3, models trained with task-aware sampling consistently outperform those trained using the full set of synthetic data that most closely resembles the training distribution. The latter is selected based on conventional similarity metrics, such as feature-wise correlation, and is often assumed to be the most representative. Our results indicate that distributional proximity alone is not a reliable proxy for downstream utility. In contrast, selecting synthetic samples based on task performance leads to more meaningful augmentation and improved predictive outcomes.

**Figure 3.**
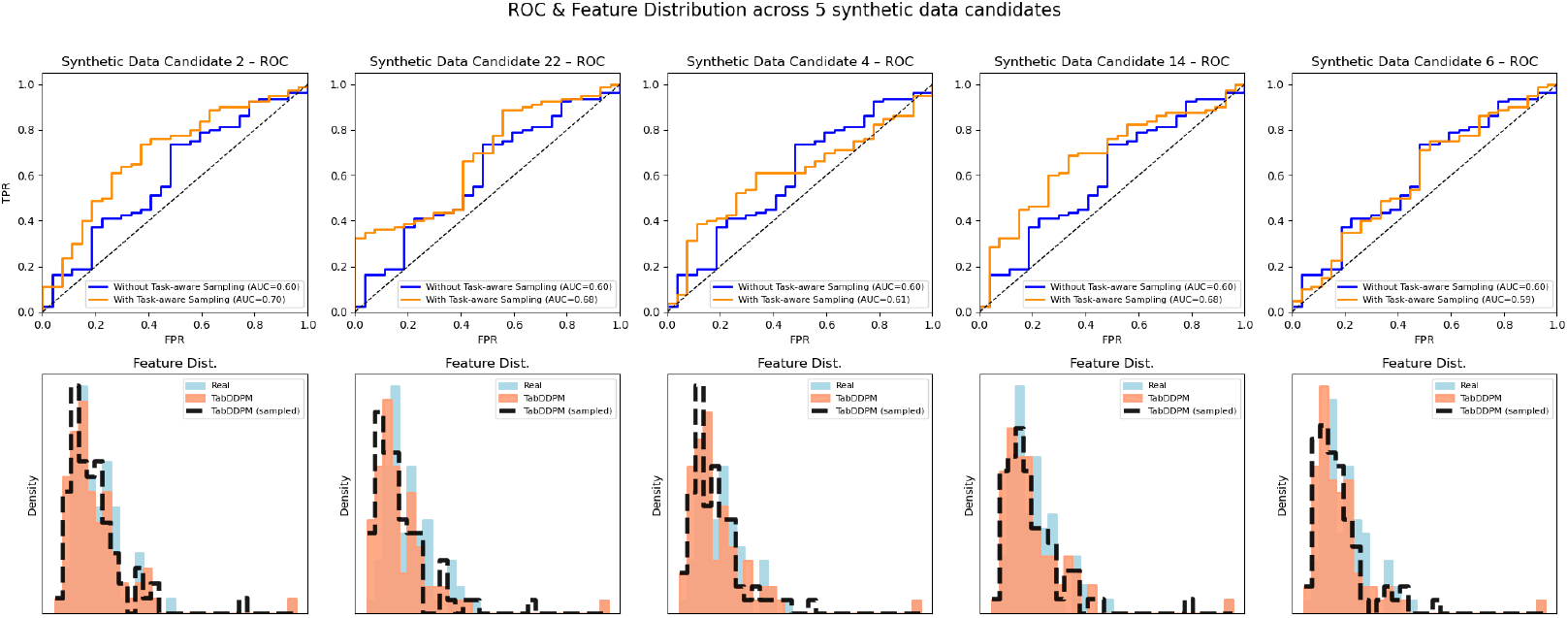
Task-aware sampling improves downstream performance. Top: ROC curves for five synthetic data candidates (orange: with task-aware sampling; blue: without task-aware sampling), which were selected based on minimal average correlation differences to real data. Although task-aware sampling introduces slight distributional shifts away from real data, it consistently improves AUC, demonstrating the benefit of targeted sampling for predictive performance. Bottom: Average feature distributions for real data (blue), the complete synthetic dataset (red), and the subset selected by task-aware sampling (black dashed). Task-aware sampling slightly alters synthetic feature distributions, enhancing diversity and providing additional information that improves final model performance.

### Domain Adaptation Enhances Cross-Cohort Generalization

While synthetic data augmentation and sampling improve data efficiency, cohort-level heterogeneity remains a key barrier to generalization in biomedical applications. Differences in patient populations, measurement protocols, or institutional biases can cause significant distribution shifts across datasets. To address this, we integrate a domain adaptation module into our pipeline, based on the Multi-Domain Adversarial Network (MDAN), which explicitly learns domain-invariant representations across real and synthetic cohorts.

In our multi-source domain adaptation network (MDAN) implementation, it aims to learn domain-invariant feature representations across heterogeneous biomedical cohorts. The architecture includes three main components: a shared *feature extractor*, a *label predictor* for the primary prediction task, and a *domain classifier* trained adversarially to encourage alignment between domains.

During training, each input sample, whether from a source or target domain, is passed through the shared *feature extractor*. Samples from labeled source cohorts are supervised using the *label predictor*, which outputs class logits and is optimized using cross-entropy loss. In parallel, all samples—including both labeled source and unlabeled target domains—are fed into the *domain classifier*, which attempts to distinguish the domain of samples. To encourage the feature extractor to generate domain-invariant representations, it utilizes a *Gradient Reversal Layer (GRL)* between the feature extractor and the domain classifier. GRL multiplies the gradient by a negative scaling factor −*α* during backpropagation (see Method Section A), effectively reversing the optimization signal and forcing the feature extractor to confuse the domain classifier. This adversarial training reduces inter-cohort discrepancies and promotes alignment across real, synthetic, and target domains.

As shown in Figure 4, incorporating a domain adaptation mechanism on heterogeneous biomedical datasets yields comparable or better AUROC compared to state-of-the-art tabular learning methods. Specifically, MDAN achieves a relative AUROC improvement of +10.3% over tree-based models (0.591 vs. 0.536), +3.7% over deep learning models (0.591 vs. 0.570), and +3.5% over TabPFN (0.591 vs. 0.571). While AutoGluon demonstrates slightly higher AUROC (0.626 vs. 0.591), it requires significantly longer training times than the MDAN network. Unlike these one-size-fits-all architectures, which implicitly assume homogeneous data distributions, MDAN explicitly models cross-cohort shifts and learns domain-invariant features. This leads to improved accuracy and substantially reduces performance gaps across heterogeneous evaluation domains.

**Figure 4.**
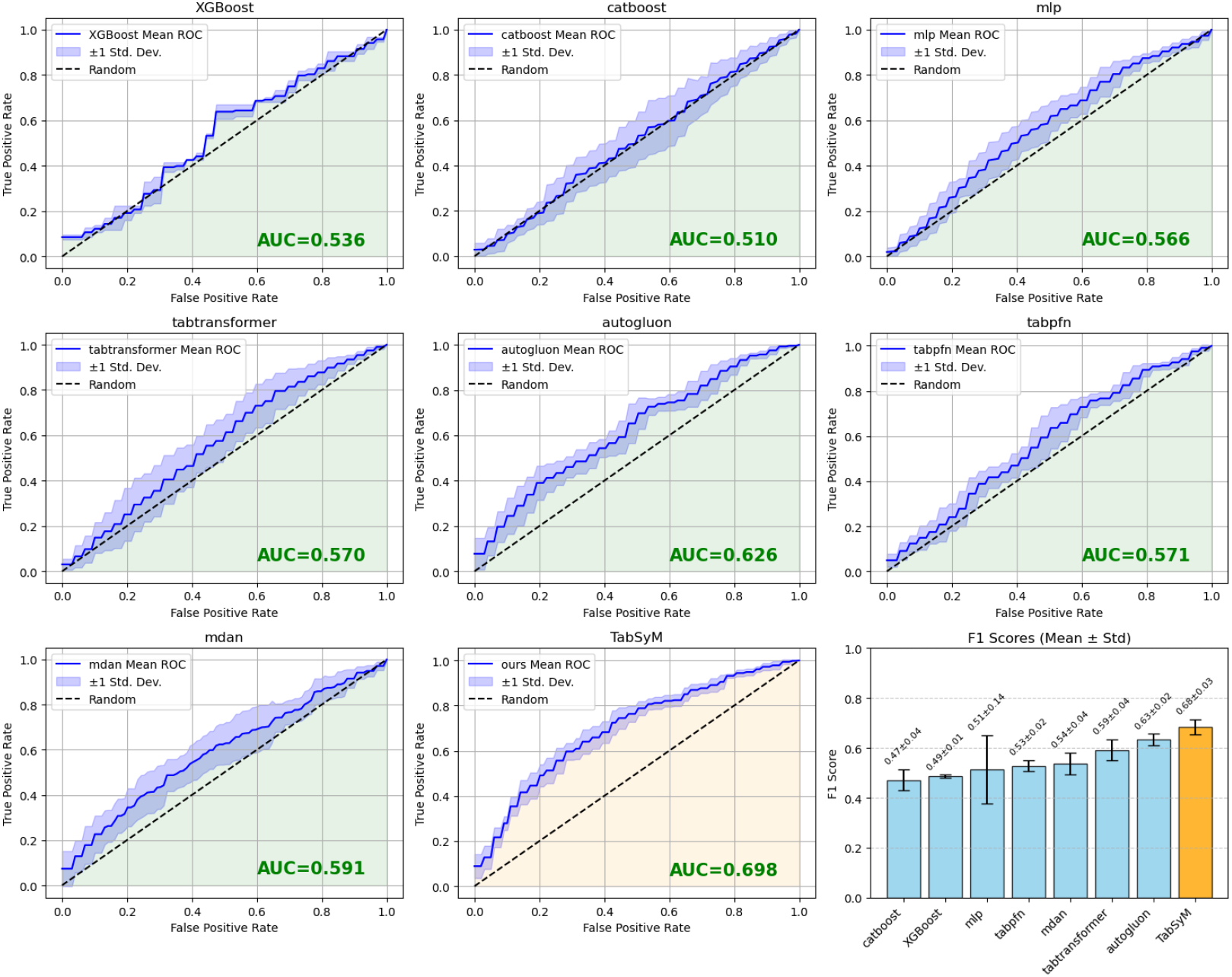
Performance on survival task across baselines.

### Performance on real-world biomedical task

We evaluate our full pipeline on a real-world biomedical prediction task designed to reflect the challenges of small sample size, cohort heterogeneity, and high-dimensional feature spaces.

The task focuses on predicting 3-year survival in gastric cancer patients based on bulk RNA sequencing data. Four GEO datasets—GSE15459 (Singapore), GSE26899 (KUGH), GSE26901 (KUCM) and GSE62254 (ACRG)—serve as source domains, while the TCGA Stomach Adenocarcinoma PanCancer Atlas cohort acts as the target domain (see Methods for details). To simulate a realistic deployment scenario, we assume limited label access to TCGA during training. Our model is trained using the four source domains, augmented with synthetic data, and incorporates multi-domain adaptation to address cohort shifts. We reserve 20% of TCGA as a held-out validation set, used exclusively for hyperparameter tuning and task-aware sample selection. The remaining 80% data will be split into a 60% training set for unsupervised MDAN training and a 20% independent test set. Since no label information is used from these two parts, their predictions will be combined to evaluate final performance.

We benchmark our pipeline against a diverse set of widely used machine learning methods for tabular data.

The following baselines span classical algorithms, deep learning architectures, and state-of-the-art automated frameworks.

### Classical tree-based algorithms

Tree-based algorithms are widely regarded as the most effective models for tabular data due to their ability to capture complex feature interactions, naturally handle missing values, and offer strong performance on heterogeneous datasets. In this study, we select two representative and widely adopted gradient boosting methods: **XGBoost** ^54^ is a highly efficient implementation of gradient boosting that uses second-order optimization and tree pruning to accelerate training and improve accuracy. XGBoost supports sparse input handling and parallel computation, making it especially well-suited for tabular datasets. **CatBoost** ^55^ is another gradient boosting algorithm developed by Yandex, distinguished by its native support for categorical features through efficient encoding, and its use of ordered boosting to reduce overfitting. CatBoost automates feature preprocessing and is optimized for fast, robust learning on structured data.

### Deep learning architectures

While classical machine learning methods have dominated tabular data modeling, a growing body of work demonstrates that deep learning architectures can achieve comparable or even superior performance in certain settings. In this study, we include two representative neural models. **MLP (Multilayer Perceptron)** ^56^ is a classical feedforward neural network composed of multiple fully connected layers, capable of capturing non-linear relationships in tabular data. MLP serves as a standard deep learning baseline for comparison. **TabTransformer** ^57^ is a more recently developed neural architecture specifically designed for tabular data, which utilizes self-attention mechanisms to model complex feature interactions and has shown improvements over traditional MLPs in various benchmarks.

### Automated/Unified learning framework

Recent advances have introduced end-to-end solutions that allow users to bypass manual hyperparameter tuning and feature engineering, streamlining the modeling process for tabular data. In our experiments, we consider two representative frameworks: **AutoGluon** ^58^ is an automated machine learning (AutoML) framework that automatically selects, ensembles, and tunes machine learning models for tabular data, delivering strong out-of-the-box performance with minimal manual effort. **TabPFN** ^59^ is a foundation model recently proposed that enables fast and robust predictions for tabular data with minimal training. TabPFN is pre-trained on a large collection of tabular tasks, allowing it to generalize to new datasets via a single forward pass.

All the above baselines are implemented in a standard supervised learning setting: we combine the four source cohorts into a single training set, and partition the target cohort TCGA into a 20% validation set for hyperparameter tuning and an 80% test set for final performance evaluation. This setup ensures that the sample allocation and evaluation protocol are consistent with those used in our pipeline, allowing for a fair and direct comparison.

For each method, we performed 10 independent runs with different random seeds and randomized train–test splits to ensure statistical robustness. All hyperparameters were tuned using Optuna, with optimization guided by performance on a held-out validation set. All baseline models, as well as our full pipeline, were executed on a shared laboratory machine equipped with three NVIDIA RTX 4090 GPUs and 32-core CPUs. Task-aware sampling procedures, which require multiple candidate generations and validation-based selection, were parallelized across CPUs and GPUs to improve runtime efficiency.

As shown in Figure 4, our method achieves superior performance on the gastric cancer survival prediction task across all categories of baselines. Compared to classical tree-based models, our pipeline significantly improves AUROC by 30.2% (0.698 compared with 0.536) and F1 score by 40.9% (0.685 compared with 0.486) over the best of XGBoost and CatBoost. For deep learning architectures, we observe an AUROC improvement of 22.5% (0.698 compared with 0.570) relative to the better of MLP and TabTransformer. When evaluated against automated learning frameworks, including AutoGluon and TabPFN, our method achieves an AUROC gain of 11.5% (0.698 compared with 0.626) and F1 improvement of 8.2% (0.685 compared with 0.633).

These results highlight the effectiveness of combining synthetic data generation, task-aware sampling, and domain adaptation to address the challenges of high-dimensional, cohort-heterogeneous biomedical datasets.

### Modular Generative Sampling Enhances Model-Agnostic Generalization

While our full pipeline integrates generative augmentation, task-aware sampling, and domain adaptation, each of these components is designed to function independently and can be flexibly integrated into other machine learning workflows. In particular, the synthetic data generation and task-aware selection mechanisms are model-agnostic by construction. They do not rely on the architectural assumptions or training dynamics of deep neural networks and can be applied to a wide range of learning tasks. This modularity makes our pipeline not only effective for domain-adversarial settings but also broadly useful in scenarios where domain adaptation is unavailable or unnecessary.

To validate the generality of our approach, we isolate the generation and sampling modules and apply them to a baseline model—XGBoost—on the gastric cancer survival task. To ensure a fair comparison, we adopt the same experimental setup as in our main pipeline: the model is trained on four labeled source cohorts (Singapore, ACRG, KUCM, and KUGH) and evaluated on the target cohort (TCGA). The TCGA samples are split into 80% for testing and 20% for validation. The validation set is used exclusively for hyperparameter tuning and task-aware synthetic sample selection, while final evaluation is conducted on the 80% test set using AUROC and F1 score as performance metrics. As shown in Figure 5, augmenting the training data with synthetic samples selected via task-aware sampling leads to a marked improvement in predictive accuracy by nearly 100% (0.748 compared with 0.374), F1 score by 80.3% (0.641 compared to 0.355) and AUROC by 7.9% (0.560 compared to 0.519) compared to training on real data alone. This result is particularly notable given that XGBoost is a non-deep, non-domain-adaptive model, suggesting that our sampling-enhanced synthetic data retains sufficient structure and diversity to benefit classical learners. These findings support the broader claim that synthetic augmentation and task-guided sampling are not tightly coupled to any specific model. Instead, they represent a general-purpose tool that can enhance downstream learning across architectures, training regimes, and data availability conditions. By designing these components to be modular and lightweight, our framework can serve as a plug-in layer for improving tabular learning in both deep and traditional modeling pipelines.

**Figure 5.**
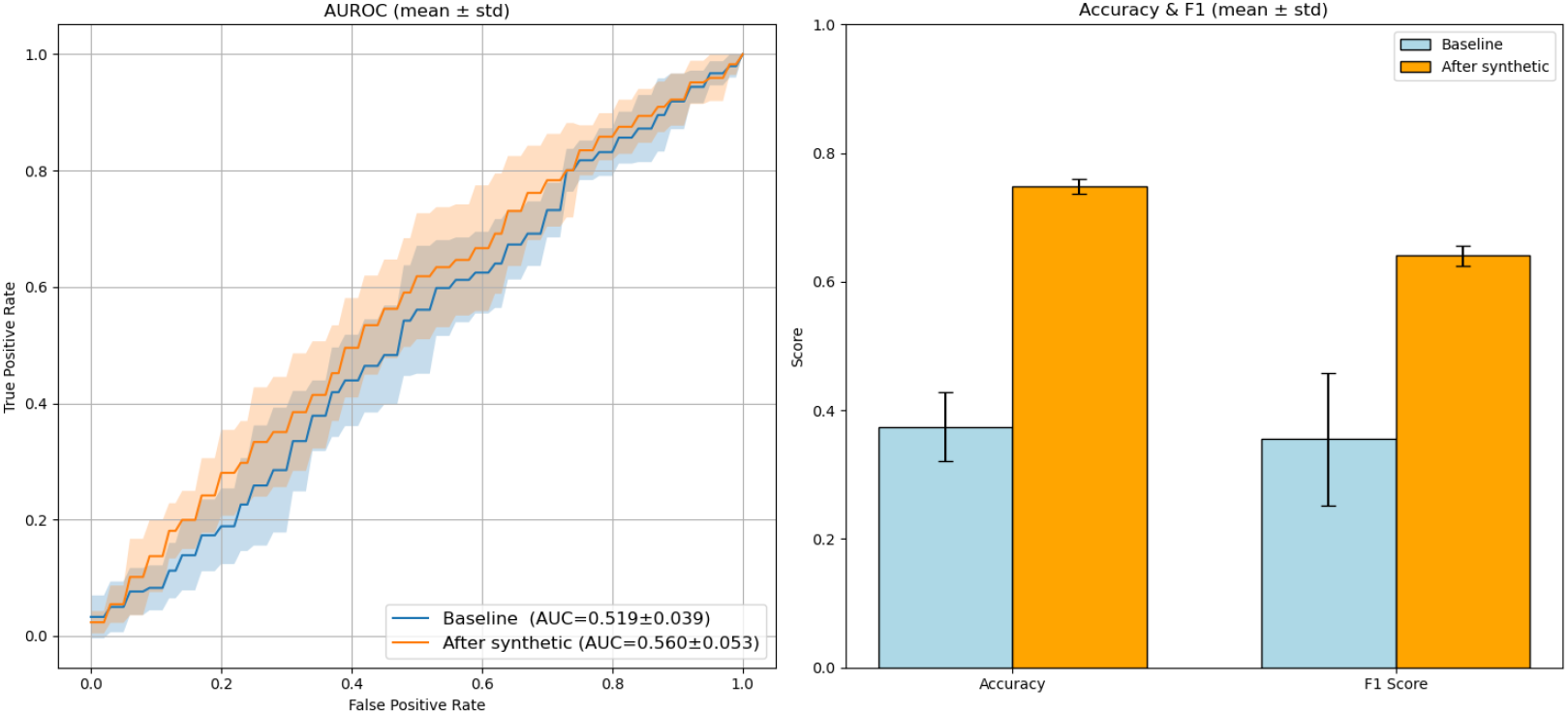
Performance on adding synthetic data to XGBoost.

## Discussion

In this work, we developed a modular learning pipeline to address the fundamental challenges of biomedical data modeling, where datasets are typically small, high-dimensional, and cohort-specific. We validated our approach on a difficult prediction task and observed significant improvements over strong baselines. Central to our method is the integration of generative modeling and task-aware sampling, which together enable more effective learning in tabular biomedical settings with limited supervision. Our experiments demonstrate that this strategy not only enhances performance on complex few-shot tasks, but also generalizes across different model families, highlighting its utility as a model-agnostic augmentation module. Altogether, our framework provides a new direction for improving learning in small-sample scenarios, especially in tabular biomedical and clinical datasets where data scarcity remains a critical bottleneck.

Despite these strengths, our current implementation faces efficiency limitations compared to state-of-the-art machine learning approaches. In particular, the need for online optimization of the generative and sampling components introduces additional computational overhead. Future work could explore ways to improve the overall runtime performance of the pipeline while maintaining its predictive benefits. Another direction is to further validate the generality of our framework by applying it to a broader range of prediction tasks and biomedical datasets. Extending the method beyond classification and evaluating it across varied cohort structures and data modalities may offer deeper insight into its practical utility across real-world settings. As machine learning continues to gain traction in the biomedical domain, where tabular data remains a dominant modality, we believe our modular synthetic learning pipeline offers a promising foundation for unlocking the full potential of data-driven modeling in this field.

## Methods

### Datasets and Problem setup

We downloaded GSE15459 (Singapore), GSE26899 (KUGH), GSE26901 (KUCM) and GSE62254 (ACRG) from the Gene Expression Omnibus (GEO). Corresponding clinical and survival information was taken from the supplementary tables of the original publications ^60 61 61 62 63^. Transcript-per-million (TPM) expression data and clinical annotations for the Stomach Adenocarcinoma PanCancer Atlas cohort (TCGA) were obtained via cBioPortal ^64^. GSE62254 was processed with Robust Multi-array Average (RMA). For the remaining GEO cohorts, we confirmed that the expression matrices were log_2_-transformed and quantile-normalised. TCGA TPM values were used as provided. For each cohort, we applied EcoTyper ^65^ in *recovery* mode with default parameters to derive 71 cell-state abundance scores per sample; these scores constitute the input features for all downstream analyses.

We consider a binary classification task for predicting 3-year survival outcomes in gastric cancer patients, where the objective is to identify whether a patient will die within three years of diagnosis. For each sample, if the recorded survival time is less than three years and the patient is deceased, we assign a positive label (1). If the patient either survived beyond three years or is deceased with a survival time exceeding three years, the label is set to negative (0). Samples for which the patient is still alive and the follow-up time is less than three years are excluded from the dataset due to the uncertainty in label assignment.

In the baseline setting without domain adaptation, we aggregate all four labeled source cohorts—Singapore (77 samples), ACRG (101 samples), KUCM (141 samples), and KUGH (300 samples)—into a single training dataset, totaling 619 samples. These are fully labeled and used for supervised training. The target cohort, TCGA, contains 179 usable samples after filtering. We split TCGA into 80% test and 20% validation. The validation set is used exclusively for hyperparameter tuning via Optuna, which is run with 20 trials for each baseline model.

For our full method and domain-adaptive variants using MDAN, we treat each source cohort as a distinct domain during training, preserving their individual distributional characteristics. Labels from each source domain are used in the supervised training phase. The target domain (TCGA) is divided into 60% unlabeled training data (used only for unsupervised domain alignment), 20% validation data (used for hyperparameter tuning and task-aware sampling), and 20% test data (used for final evaluation). The validation and test subsets of TCGA are entirely excluded from training. Final evaluation is conducted on the union of the unlabeled target training set and the held-out test set to assess generalization under realistic deployment conditions.

### Synthetic data generation

To mitigate sample scarcity in biomedical settings, our generator is adapted from the official open-source implementations of TabDDPM ^43^, a modification of DDPM ^28^ and multinomial diffusion ^66^, combining continuous and categorical modeling for tabular data. Although TabDDPM supports both numerical and categorical features, our datasets contain only continuous (numerical) features, and we therefore omit the categorical modeling components.

We train a single TabDDPM model by aggregating all four labeled source datasets (Singapore, ACRG, KUCM, KUGH), using both feature vectors and survival labels. Target domain data (TCGA) is strictly excluded from training to prevent leakage.

The TabDDPM model learns to reverse a noising process applied to the original data. Following DDPM, Gaussian noise with a cosine-based *β*-schedule 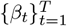 is added to each numerical vector *x*_num_. Let *α*_*t*_ = 1 − *β*_*t*_ and 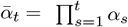. The *t*-step marginal is

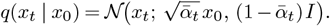

The denoising network ***ϵ***_*θ*_(*x*_*t*_, *t, y*) learns to predict either the original sample *x*_0_ or the added noise ***ϵ***, conditioned on timestep *t* and survival label *y*.

Label guidance is incorporated by conditioning the denoising model on the survival label *y*. TabDDPM utilizes a simple MLP architecture adapted from Gorishniy *et al.* ^67^ to model the reverse process:

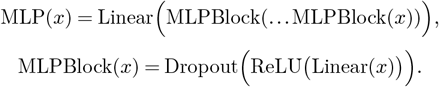

The TabDDPM implementation, following Nichol *et al.* ^68^, Dhariwal & Nichol ^69^, process a tabular input *x*_in_, a timestep *t*, and a class label *y* as follows:

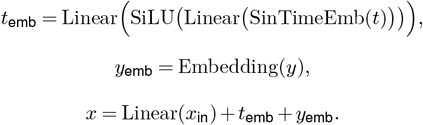

Here, SinTimeEmb denotes a sinusoidal time embedding with a dimension of 128. All Linear layers in these equations use a fixed projection dimension of 128. During training, the model minimizes the following simple denoising loss for numerical features:

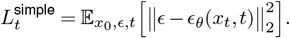

TabDDPM also handles categorical features using a KL divergence term in the loss function. However, since no categorical features are present in our dataset, the categorical component of the loss is omitted. During inference, we sample synthetic points conditioned on each class proportionally to the label distribution in the training data.

To enhance robustness and avoid mode collapse, we retain synthetic outputs from various hyperparameter settings (e.g., trained from 20 Optuna trials), increasing both variance and diversity in the generated pool. The Optuna search space is defined as follows:

- Learning rate (*η*): `loguniform`[10^−5^, 5 *×* 10^−3^]
- Model depth (number of hidden layers): {2, 3, 4}
- Batch size: {256, 4096, 8192, 16384, 32768}
- Total diffusion training steps: {5000, 20000, 30000}
- Number of diffusion timesteps: {100, 1000}

The number of generated samples is dynamically scaled based on the training set size. The objective function used in Optuna maximizes the F1 score from a pre-tuned MLP model trained on the generated data.

#### A. Multi-source domain adaptation

To align heterogeneous source and target domains, we implement a Multi-source Domain Adversarial Network ^40^ (MDAN) that learns domain-invariant representations via adversarial training. As illustrated in Figure 2, the model consists of three key components: a shared feature extractor, a label predictor, and a domain classifier.

The feature extractor is composed of residual MLP blocks with LeakyReLU activations, dropout, and batch normalization layers.

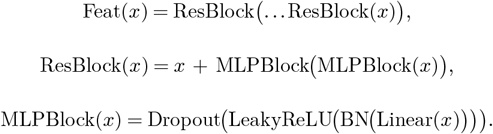

The extracted features are normalized via a LayerNorm layer, and optionally refined using a self-attention module. These domain-invariant features are simultaneously used to predict task labels and domain identities.

The label predictor is a two-layer MLP trained in a supervised fashion on labeled source and synthetic data. It is optimized using a standard cross-entropy loss.

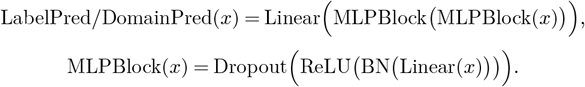

The domain classifier is also a two-layer MLP, but receives input features passed through a Gradient Reversal Layer (GRL).

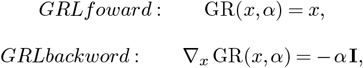

The GRL multiplies the gradients from the domain classifier by −*α* during backpropagation, effectively encouraging the feature extractor to produce features that are indistinguishable across domains. The domain classifier is optimized with a domain classification loss *ℒ*_dom_, using cross-entropy to predict the domain label (e.g., source real, source synthetic, or target). The target domain samples are included without labels but are used for domain discrimination.

The overall multi-domain adversarial network (MDAN) produces label-specific and domain-specific logits in a single forward pass:

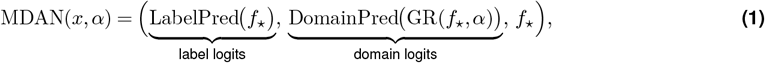

The model is trained by jointly minimizing a task-classification loss ℒ_label_ and an adversarial domain loss ℒ_dom_:

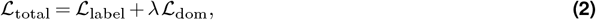

where the hyper-parameter *λ >* 0 controls the strength of the domain alignment term,which is also tuned using optuna. Minimizing ℒ_label_ encourages accurate label prediction, whereas the gradient-reversed ℒ_dom_ drives the feature extractor to learn domain-invariant representations.

All source domains (Singapore, ACRG, KUCM, and KUGH) and their corresponding synthetic samples are used with full label supervision. The target domain (TCGA) is included only for domain alignment without using its label information. This allows our framework to generalize to unlabeled cohorts through adversarial feature alignment.

We use Optuna to tune the MDAN hyperparameters. The objective is to maximize AUROC on a held-out validation set from the target domain. We search over:

- Learning rate (*η*): `loguniform`[10^−5^, 5 *×* 10^−3^]
- Hidden layer sizes: First layer ∈ 64, 96,*…,* 256; second layer ∈ 32, 64, 96, 128
- Dropout rate: `uniform`[0.2, 0.6]
- Domain adversarial weight (*λ*): `uniform`[0.1, 2.0]
- Weight decay: `loguniform`[10^−6^, 10^−2^]
- Mixup alpha (for regularization): `uniform`[0.1, 0.8]
- Use of attention: `True`or `False`

We run 20 Optuna trials and train for up to 300 epochs with early stopping (patience = 15). All models are trained on a shared lab machine with three NVIDIA RTX 4090 GPUs and 32 CPU cores.

#### B. Task-Aware Sampling Strategy

To effectively utilize synthetic data while avoiding overfitting or noise amplification, we incorporate a task-aware sampling module that automatically selects the most informative synthetic subsets. This module operates on the pool of 20 synthetic datasets generated by TabDDPM under different hyperparameter configurations (selected via Optuna as described in Section), with each synthetic group reflecting a unique distribution.

For each candidate group, we formulate the sampling process as a black-box optimization problem over two continuous variables: the sampling proportions *p*_0_ and *p*_1_ for class 0 and class 1, respectively. These parameters determine the fraction of synthetic samples per class to be included in the training process. The optimization is conducted using Gaussian Process-based Bayesian Optimization, implemented via the `skopt`package. The search space is *p*_0_, *p*_1_ ∈ [0.0, 1.0], with a total of 20 Bayesian Optimization evaluations (5 initial points and 15 model-guided iterations). To ensure better initialization and coverage of the sampling space, we fix the following initial points:

~~~
custom_x0 = [
  [1, 1],
  [1, 0.5],
  [0.5, 1],
  [0.5, 0.5],
  [0, 0.5],
  [0.5, 0],
]
~~~

For each (*p*_0_, *p*_1_) candidate, the corresponding subset is sampled from the synthetic group, appended to the list of source data, and used to train an MDAN model with fixed architecture and best-tuned hyperparameters. Validation AUROC on a held-out portion of the target domain (TCGA) is used as the Bayesian Optimization objective function. Importantly, this validation set (20% of TCGA, disjoint from training and test sets) is used exclusively for sampling and hyperparameter selection, not for final evaluation.

To improve robustness, we perform Bayesian Optimization independently for all 20 synthetic groups in parallel, and retain the top-5 performing subsets ranked by validation AUROC. At inference time, prediction scores from models trained on each of these five selected configurations are averaged to produce the final output.

All models are trained using the same MDAN architecture as described in Section A, and the sampling pipeline is implemented in Python with multiprocessing to accelerate Bayesian Optimization execution across synthetic groups. Each model uses the same data split and shares validation and test configurations for consistency.

